# A New European Species of The Opportunistic Pathogenic Genus *Saksenaea* – *S. dorisiae* Sp. Nov.

**DOI:** 10.1101/597955

**Authors:** Roman Labuda, Andreas Bernreiter, Doris Hochenauer, Christoph Schüller, Alena Kubátová, Joseph Strauss, Martin Wagner

## Abstract

A new species *Saksenaea dorisiae* (Mucoromycotina, Mucorales), recently isolated from a water sample originating from a private well in a rural area of Serbia (Europe), is described and illustrated. The new taxon is well supported by phylogenetic analysis of the internal transcribed spacer region (ITS), domains D1 and D2 of the 28S rRNA gene (LSU), and translation elongation factor-1α gene (TEF-1α), and it is resolved in a clade with *S. oblongispora* and *S. trapezispora*. This fungus is characteristic by its moderately slow growth at 15 and 37°C, sparse rhizoids, conical-shaped sporangia and short-cylindrical sporangiospores. *S. dorisiae* is a member of the opportunistic pathogenic genus often involved in severe human and animal mucormycoses encountered in tropical and subtropical regions. Despite its sensitivity to several conventional antifungals (terbinafine and ciclopirox), the fungus is potentially causing clinically challenging infections. This is the first novel taxon of the genus *Saksenaea* described from the moderately continental climate area of Europe.

## INTRODUCTION

The genus *Saksenaea* S.B. Saksena belongs to mucoralean microscopic fungi (Mucoromycotina, Mucorales, Saksenaeaceae) comprising species often causing severe human and animal cutaneous mucormycoses in both immunocompromised and immunocompetent hosts (1, 2). The genus was described from a forest soil in India in 1953, with *Saksenaea vasiformis* S.B. Saksena being a type and for more than 50 years a single-species of the genus (nowadays considered as *S. vasiformis* species complex), reported in soil, drift-wood, and grains (1). Interruption of skin integrity, such as needle sticks, insect stings or spider bites, burn and accident wounds (up to 95% of cases) represent the most common mode of infection by this fungus from contaminated soil or water (2–4). In addition to these traumatic implantations infections through inhalation of spores and the use of indwelling catheters have been reported (5). The species of the genus are responsible for skin infections, characterized by rapid progression and invasion of neighboring tissues. Infection of the *Saksenaea* fungi is diagnosed by angio-invasion leading to tissue necrosis with cutaneous and subcutaneous involvement. Furthermore, rhino-orbito-cerebral and disseminated infections have been reported (1, 2). Classic management of the infection site usually includes a combination of broad, aggressive and repeated surgical debridement (which may lead to amputation), and long–term systemic therapy with appropriate antifungals, preferably liposomal amphotericin B and/or posaconazole (6–9). Antifungal therapy alone seems to be inadequate to control infection (2).

Recent revisions by Alvarez et al. (1) and Crous et al. (10, 11) applying a polyphasic approach including sequence analysis of three loci (ITS, LSU, EF-1α) revealed that a monospecific genus *Saksenaea* is actually genetically heterogenous and encompasses more species, namely *S. vasiformis* complex (with more putative cryptic species), *S. loutrophoriformis*, *S. erythrospora*, *S. oblongispora* and *S. trapezispora*. So far these species (except *S. oblongispora*) have been reported worldwide from human and animal clinical cases. They were encountered in tropical and subtropical climates and have been reported from Australia, Thailand, India and Sri Lanka, the Middle East, Tunisia, the USA, Central and South America (1, 12). In Europe, there are only a few published cases available reporting *S. vasiformis* infections, e.g. from Spain (3, 13–15), France (1), and Greece (2). Up to now, approximately 45 cases of mostly cutaneous infections of *Saksenaea*, have been reported around the world (2, 4, 5, 10, 11), although the actual number of clinical cases is possibly underestimated (4). Despite its reproducibility in severe human infections, this genus remains a poorly studied mucoralean genus, mainly due to the lack of sporulation on the mycological culture media (e.g. sabourad agar, malt extract agar, corn-meal agar) routinely used in clinical laboratories (1).

During our microbiological survey of water samples a fast growing mucoralean nonsporulating fungus was recovered on a *Pseudomonas* isolation plate in a sample originating from a private well in a rural area of Manastirica (Serbia) in October 2018. This isolate was designated BiMM-F232 and further characterized in terms of morphology, physiology, molecular phylogeny and antifungal susceptibility. Phylogenetically informative sequences were obtained from three loci, i.e. internal transcribed spacer region including 5.8S rDNA (ITS), domains D1 and D2 of the 28S rRNA gene (LSU), and from the translation elongation factor-1α locus (TEF1α). Overall, the resulting data revealed that this isolate represents a novel species of the opportunistic pathogenic genus *Saksenaea*, and it is described and illustrated here as *Saksenaea dorisiae* sp. nov.

## MATERIALS AND METHODS

### Sample collection and isolation of the fungus

A single sample of water from a private well in Manastirica-Petrovac (The Republic of Serbia, Europe) was collected in October 2018. A 100 mL aliquot was aseptically filtered through a filter paper disc (0.45 µm, 47 mm diameter, GN-6Metricel, Pall, Mexico) plated on *Pseudomonas* Selective Agar amended with Glycerol (Carl Roth, Germany) and incubated for 3 days at 37 (±0,2°C) in the dark.

### Cultivation of a strain, media and morphological analysis

For phenotypic determination, the strain was transferred (one agar block grown on CYA, ca 5 × 5 mm, in the middle) on Potato Dextrose Agar (PDA, Fluka), Malt Extract Agar (MEA, Merck), Corn Meal Agar (Oxoid), Water Agar 1% (WA), Oatmeal Agar (OA), Synthetic Nutrient-poor Agar (SNA), Czapek Agar (CZA), Czapek Yeast extract Agar (CYA) and Yeast Extract Sucrose Agar (YES) as described by Samson et al. (16), and incubated for 5-30 days in the dark at 25°C. Colony size (in mm), colony structure and characteristics were noted after 4 days, however the cultivation was prolonged up to 3 months in order to observe and record changes in pigmentation of the colonies as well as to determine the onset of sporangia and zygospore formation. For sporangial development, a recommended method of floating agar blocks in yeast extract water according to Padhye et Ajello (17) was used in this study. In order to determine the optimal and minimal/maximal temperatures for growth, the strain was incubated on four different media (CZA, CYA, PDA, and MEA) at 12, 15, 20, 25, 30, 35, 37, 39, 40, and 42 °C (± 0.1-0.2°C). Colony diameters were measured on the 4^th^ day of cultivation. For comparative description of the macroscopic and microscopic characteristics, CZA was used according to Alvarez et al. (1) and Crous et al., (10, 11).

The capability of three different vegetative form of the fungus to germinate or grow on more extreme temperatures was assessed by incubation at 5±0.3°C during 1 to 50 days or at 40±0.1°C for 24 to 48 hours. For these tests the following procedure was applied: (a) spores-100 µL of spore suspension (4.0 × 10^5^ CFU/mL physiologic solution) collected from CZA (6 days at 30°C) was applied on MEA plate (at 37°C) at defined time intervals (1, 2, 3, 4, 7, 14, 21, 28, 42 and 50 d); (b) mycelial pellets-micro-colonies of ca 50-200 µm in diam. (~ 2.0 × 10^4^/mL) after 10-12 d incubation submerse YES5%, 140 rpm, 25°C, 100 µL of pellet suspension applied on MEA plate (at 37°C); (c) mycelium – as a nonsporulating colony growing on MEA plate after 4 d incubation (37°C). Both spores and pellets were exposed to 5°C in physiological or YES broth liquids, respectively, and then transferred (100 µl onto a plate at 37°C). For exposure to 40°C, they were directly applied onto plates and after 24 and 48 hours transferred on a new MEA plate and further incubated at 37°C. Germination of spores was determined *in situ* under low magnification (50 –100×).

A dried herbarium specimen of the holotype was deposited in the herbarium of the Mycological Department, National Museum in Prague, Czech Republic (PRM); the ex-type cultures were deposited in the Bioactive Microbial Metabolites (BiMM) Fungal Collection, UFT-Tulln (AT) and in the Culture Collection of Fungi (CCF), Prague (CZ).

For determination of microscopic traits, CZA was used after 6-14 days. Sporangiophore structures and sporangia formation were observed *in situ* under low magnification (50 – 100×). Details in sporangiophores, sporangia, sporangiospores and other microscopic structures, such as width of hyphae, were observed in mounts with lactophenol blue (RAL Diagnostics). Sporangiospores were measured in water to prevent their deformation. Except on CZA, formation of sporangia was not observed, in any of the other media used after 8 days of cultivation. For these media incubation was prolonged for 3 months and the plates were checked at 5-day intervals for the onset of sporangia production. The photomicrographs were taken using a Motic BA 310 microscope with Motic Image Plus 3.0 software. Lactophenol blue was used as a mounting medium for microphotography. Photographs of the colonies were taken with a Sony DSC-RX100.

### Antifungal susceptibility testing

A 50 µL aliquot of sporangiospore suspension (4.0 × 10^5^ CFU/mL) of *Saksenaea dorisii* strain BiMM-F232 (collected from CZA grown at 30°C for 14 days into physiological solution) was evenly distributed on RPMI-1640 (Sigma) medium plates. In addition to the spores, a 50 µL aliquot of suspension of mycelial pellets (micro-colonies sized roughly 50-200 µm in diameter collected from YES 5% liquid medium grown at 25°C, 150 rpm after 10 days) was inoculated at a density of 2.0 × 10^4^/mL in the same way on the RPMI plates. Antifungal susceptibility testing was performed by applying the antifungal discs with 10 and 15 µg amphotericin B, 10 µg griseofulvin, 5 µg caspofungin, 1 µg flucytosin, 100 IU nystatin, 10 µg econazole, 25 and 100 µg fluconazole, 1 µg voriconazole, 10 and 15 µg ketoconazole, 5 µg posaconazole, 10 µg miconazole, 8 and 50 µg itraconazole, and 50 µg clotrimazole (Antifungal Discs, Liofilchem Diagnostici, Via Scozia, Italy), 50 µg ciclopirox and 30 µg terbinafine (Neo-Sensitabs, Rosco Diagnostica, Taastrup, Denmark) directly on the plates. MIC values were measured by applying MIC test strip with amphotericin B, flucytosin, caspofungin, ketoconazole, posaconazole, itraconazole, and voriconazole with a concentration range of 0.002-32 µg, and fluconazole with a concentration range of 0.016-256 µg (Antifungal Discs, Liofilchem Diagnostici, Via Scozia, Italy). A single MIC test strip was applied per plate, while 4-6 discs per a plate and incubated for 3-7 days at 37°C.

### DNA extraction, PCR amplification and sequencing

DNA was extracted using a standard cetyltrimethyl ammonium bromide (CTAB) procedure, as described previously (18). Polymerase chain reaction (PCR) analysis was performed by amplification of the internal transcribed spacer (ITS) region with primers ITS1-F (19) and ITS4 (20). Partial translation elongation factor (TEF-1α) gene was amplified with primers 983F (21) and 2218R (22). The D1*/*D2 domains of the large-subunit (28S) rRNA gene were amplified and sequenced using the primer pair ITS1/TW14 (20, 23). All reactions were performed in an Eppendorf Gradient MasterCycler (Eppendorf, Hamburg). Conditions for amplification of ITS and D1/D2 domains: 95°C for 5 min; 35 cycles of 95°C for 30 s, 54°C for 30 s, 72°C for 90 s and finally 5 min at 72°C. Touchdown amplification of TEF was performed as follows: 95°C for 5 min; 9 cycles of 30 s at 95°C, 30 s at 66°C (−1°C every cycle) and 1 min at 72°C, followed by 30 cycles of 30 s at 95°C, 30 s at 56°C and 1 min at 72°C and a final elongation step of 7 min at 72°C.

The PCR products were sequenced with the same primers used for the PCR amplifications (Microsynth AG, Balgach, Switzerland). All sequences obtained in this study were deposited in GenBank. For information on fungal strains used in this study see Table 1. The list provides GenBank accession numbers to ITS, TEF-1α and D1/D2 domains of 28S rRNA gene (LSU) sequences for all accepted species in the genus *Saksenaea*.

**TABLE 1.**
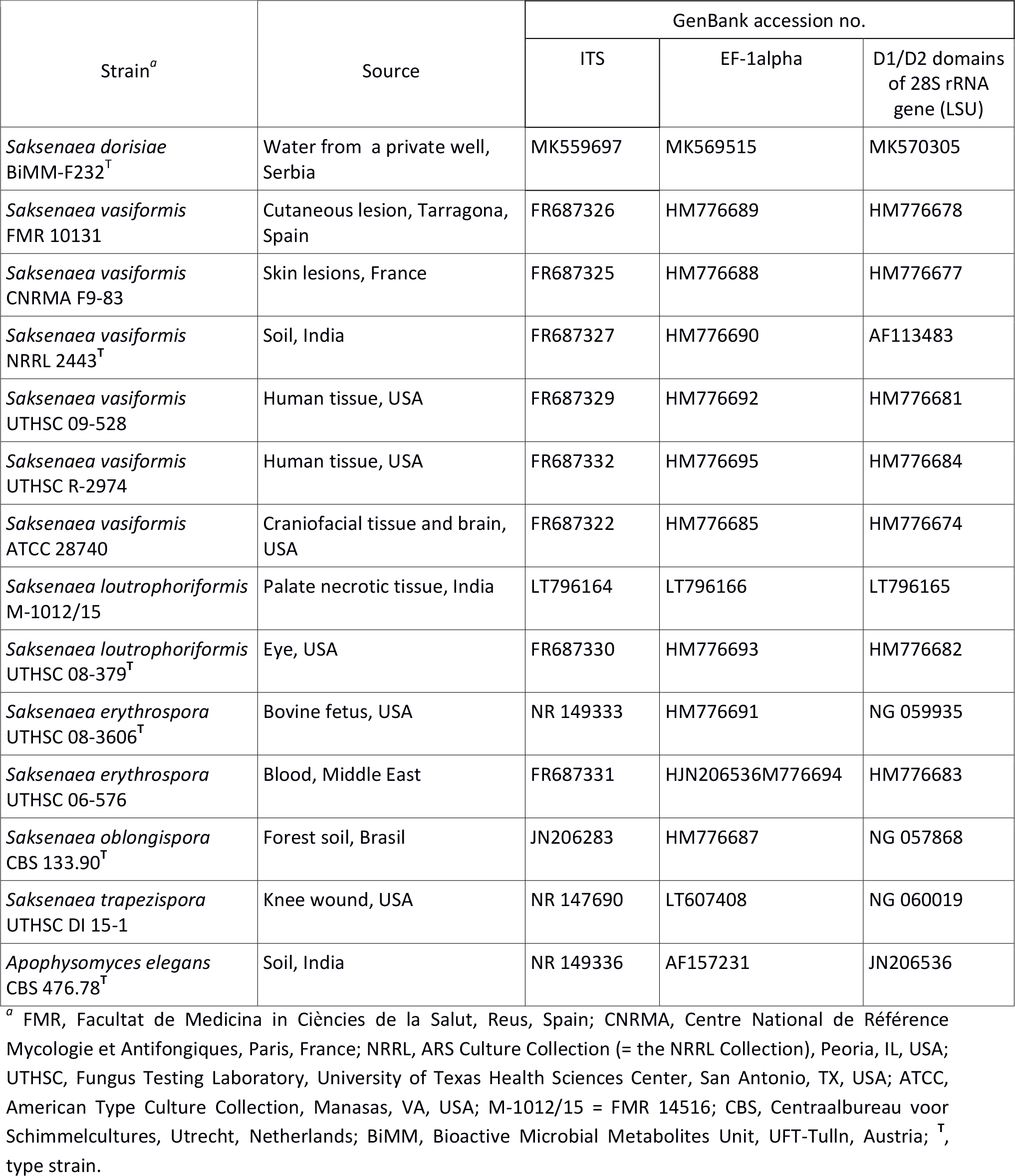
List of strains included in the study

**TABLE 2.** Comparison of the main phenotypic characteristics of the species in the genus *Saksenaea*.

| Species | Growth at 37°C on CZA after 4d (mm) | Growth at 42°C | Sporangiophore length (µm) | Shape of mature sporangia-venters ** | Spore size (µm) | Spore shape | References |
| --- | --- | --- | --- | --- | --- | --- | --- |
| *S. vasiformis* complex | >85* | + | 65-100 | spherical | 5-7 x 2-3 | long cylindrical | 1 |
| *S. loutrophoriformis* | >85 | + | 50-75 | spherical | 3.5-7x2-3.5 | long cylindrical | 11 |
| *S. erythrospora* | >85 | + | 100-150 | spherical | 5-5.5x2.5-3 | ellipsoid-biconcave | 1 |
| *S. oblongispora* | >85 | - | 80- 100 | spherical | 5-6.5x3-4.5 | oblong | 1 |
| *S. trapezispora* | 35-40 | - | 150-230 | spherical | 5.5-8x3.5-4 | trapezoid | 10 |
| ***S. dorisiae* sp. nov.** | **25-30** | - | **75-130 µm** | **conical** | **4.5-6 x2.5-3** | **Short cylindrical - capsulate** | **This study** |

\*Filling the Petri dish (diameter, 9 cm); \*\* shape of sporangia listed are based on available drawings and illustrations in the referred studies

### Phylogenetic analysis

For phylogenetic analysis, sequences were aligned with MUSCLE algorithm (24) implemented in SEAVIEW 4.6 (25) software. Phylogenetic trees were constructed using Neighbor Joining (NJ) method in SEAVIEW and genetic distances were computed with the Kimura‐2‐parameter (K2P) model. Bootstrap analyses were performed in NJ with 1000 bootstrap replicates. *Apophysomyces elegans* CBS 476.78^T^ was selected as outgroup for phylogenetic evaluation.

## RESULTS

### Taxonomy

#### Saksenaea dorisiae

R. Labuda, A. Bernreiter, C. Schüller, J. Strauss & M. Wagner **sp. nov.** Figs 1, 2 and 3

**FIG 1.**
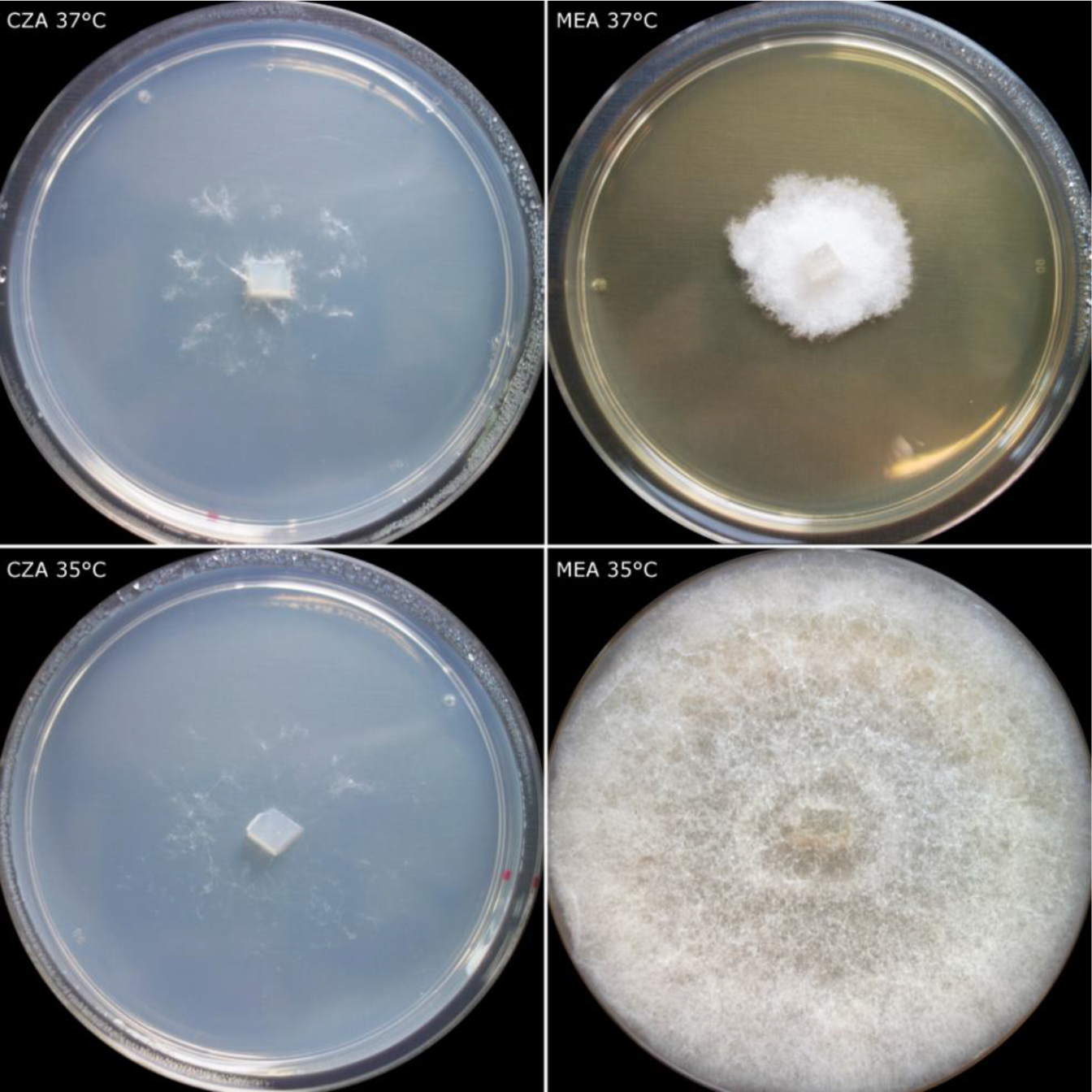
*Saksenaea dorisiae* (BiMM-F232). Colonies on CZA and MEA (4 days old) at 35 (bottom) and 37°C (top)

**FIG 2.**
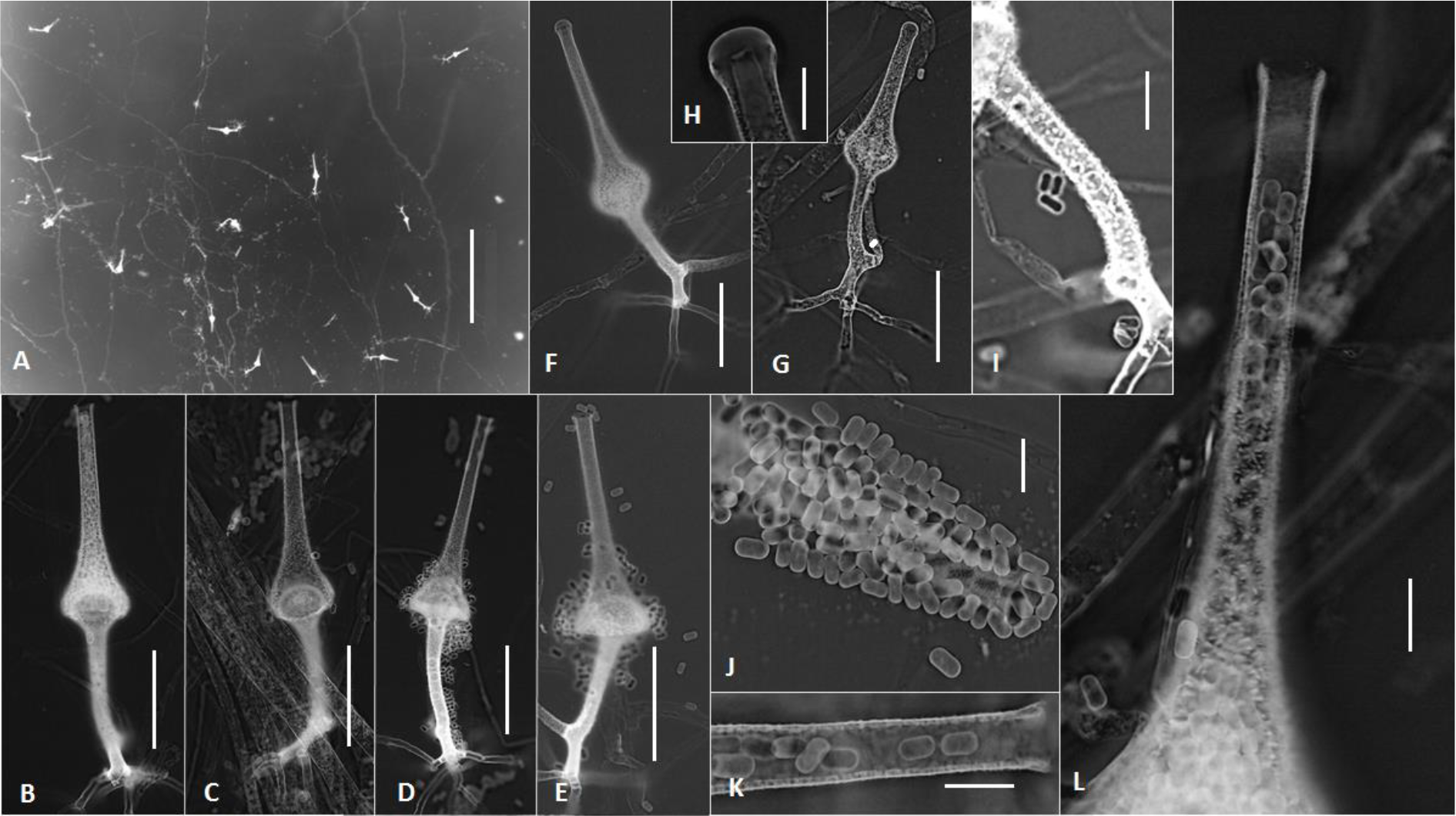
*Saksenaea dorisiae* (BiMM-F232). (A, B, C, D and E) sporangiophores with sporangia (on CZA, 6 days old). (F, G and H) young sporangia with rounded neck (closed). (I) detail of asperulate sporangiophore on CZA (6 days old). (J, K and L) sporangiospores and details of sporangial neck (on CZA, 6 days old). Scale bars = 500 µm (A), 50 µm (B-G), 10 µm (H-L).

**FIG 3.**
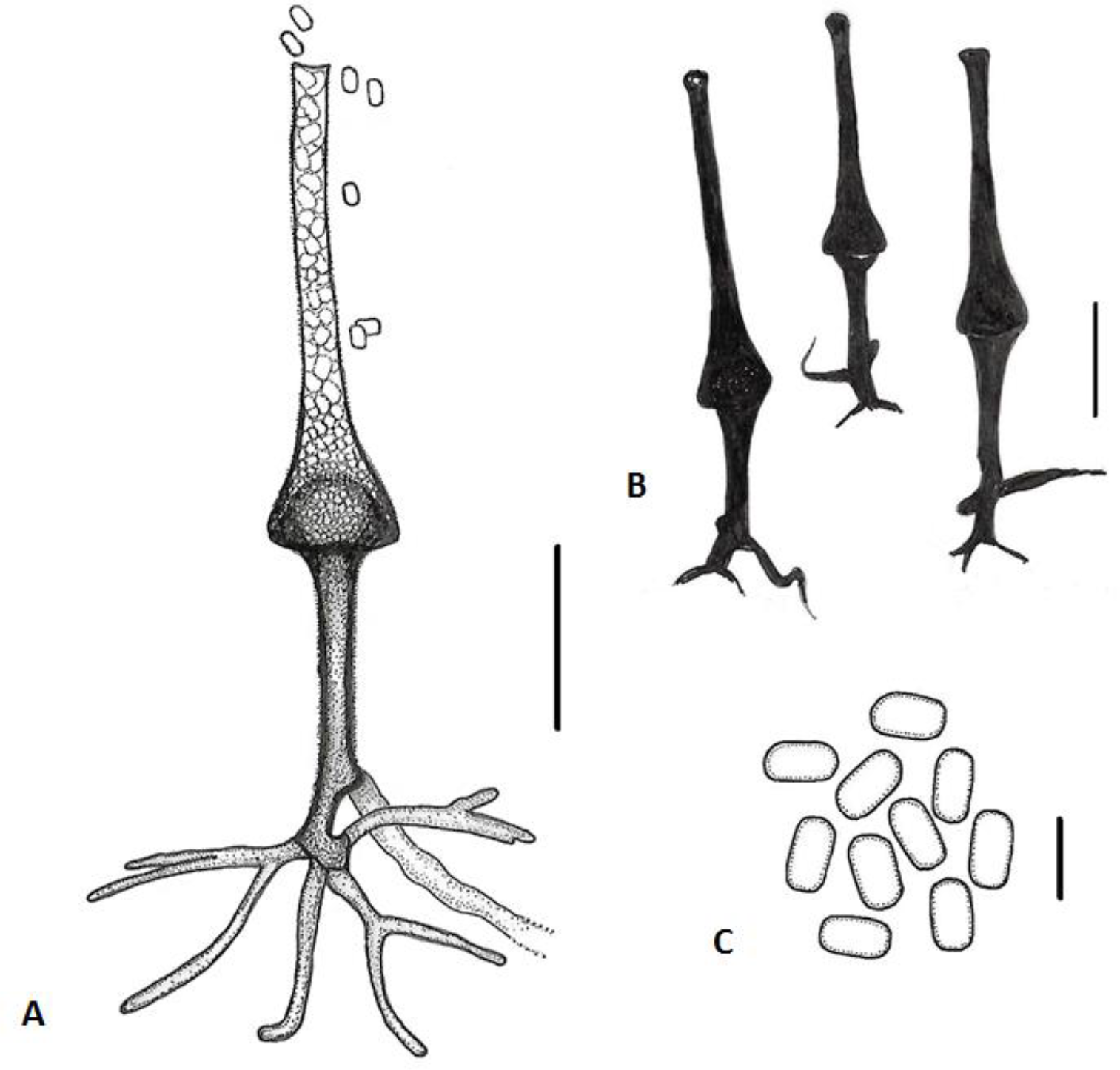
Line drawing of micromorphology of *Saksenaea dorisiae* (BiMM-F232). (A, B and C) sporangiophores, sporangia and sporangiospores on CZA (6-8 days old). (A) sporangiophore with sporangium and mature sporangiospores. (B) sporangiophores with sporangia (*in situ*). (C) sporangiospores. Scale bars = 50 µm (A-B), = 5 µm (C).

MycoBank MB 830072

*Etymology:* Latin, *dorisiae =* named after Doris Hochenauer, who isolated the fungus.

Culture characteristics (Fig. 1) – Colonies reaching 25-30 mm in diam. after 4 d of incubation at 37°C on CZA, whitish, with very scare (cobweb-like) aerial mycelium, with colorless reverse. Colonies at 37°C on MEA, CYA, PDA, CMA and SAB more abundant floccose, mycelium moderately slow growing (30-65 mm) after 4 days, with colorless reverse, remaining sterile (without sporulation).

The optimum temperature for growth on CZA was between 20 and 35 °C (45-60 mm diam.), reduced growth was observed at 15°C (15-20 mm diam.) and 37°C (25-30 mm diam.). Minimum growth was observed at 12°C (3-4 mm diam.), and the maximum temperature for growth was 39°C (0.5-1 mm diam.). The fungus did not grow at 40°C on any of the media used (CZA, CYA, MEA and PDA). An overall growth on CYA, MEA and PDA was approximately 20-25 % faster after 4 days of incubation at the optimum temperatures compared to CZA (Fig. 4). Germination and growing ability of the spores, mycelial pellets and mycelium, after exposure of 5°C and 40°C is listed in Table 3 and 4.

**TABLE 3.**
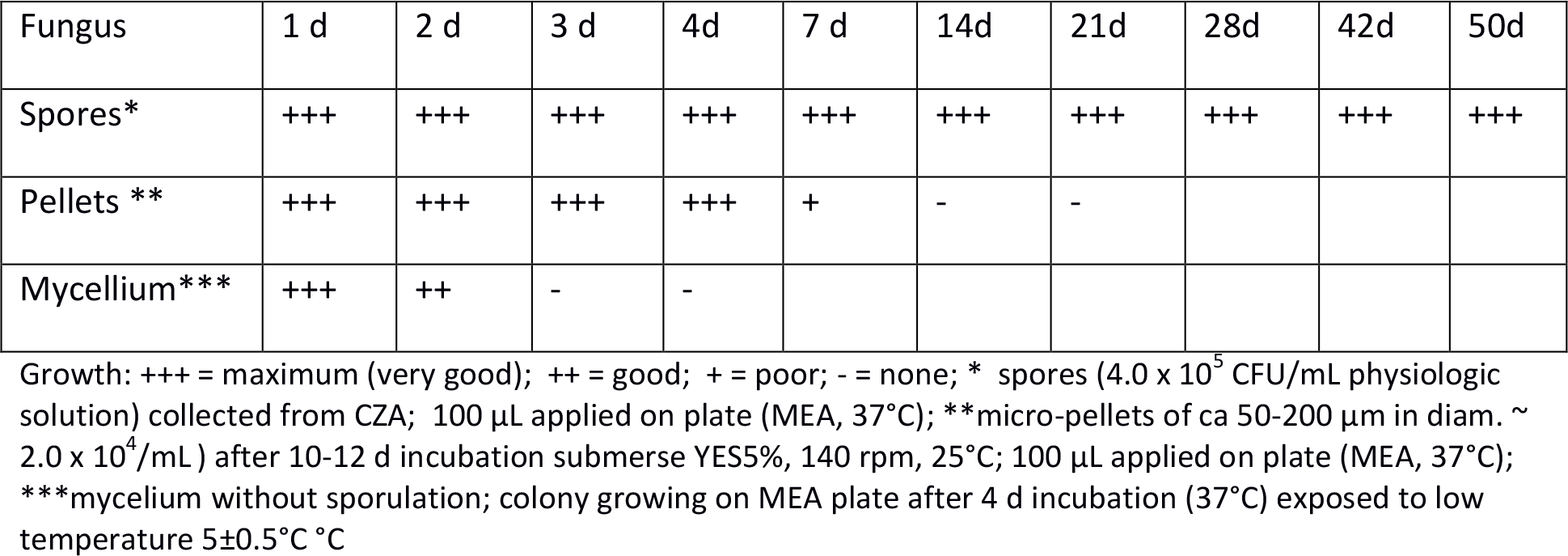
Effect of low-temperature (5±0.2°C) on viability of *Saksenaea dorisi*ae (BiMM-F232)

| Fungus | 1 d | 2 d | 3 d | 4d | 7 d | 14d | 21d | 28d | 42d | 50d |
| --- | --- | --- | --- | --- | --- | --- | --- | --- | --- | --- |
| Spores* | +++ | +++ | +++ | +++ | +++ | +++ | +++ | +++ | +++ | +++ |
| Pellets ** | +++ | +++ | +++ | +++ | + | - | - |  |  |  |
| Mycellium*** | +++ | ++ | - | - |  |  |  |  |  |  |

Growth: +++ = maximum (very good); ++ = good; + = poor; - = none; \* spores ( $4.0 \times 10^5$ CFU/mL physiologic solution) collected from CZA; 100 $\mu\text{L}$ applied on plate (MEA, $37^{\circ}\text{C}$ ); \*\*micro-pellets of ca 50-200 $\mu\text{m}$ in diam. $\sim 2.0 \times 10^4/\text{mL}$ ) after 10-12 d incubation submerge YES5%, 140 rpm, $25^{\circ}\text{C}$ ; 100 $\mu\text{L}$ applied on plate (MEA, $37^{\circ}\text{C}$ ); \*\*\*mycelium without sporulation; colony growing on MEA plate after 4 d incubation ( $37^{\circ}\text{C}$ ) exposed to low temperature $5\pm0.5^{\circ}\text{C}$

**TABLE 4.** Effect of high-temperature (40 ±0.1°C) on viability of *Saksenaea dorisiae* (BiMM-F232)

| Fungus | $40^\circ\text{C}$ (24h) | $40^\circ\text{C}$ (48h) |
| --- | --- | --- |
| Spores* | - | - |
| Pellets** | - | - |
| Mycelium*** | + | - |

Growth: + = poor; - = none; \* spores ( $4.0 \times 10^5$ CFU/mL physiologic solution) collected from CZA (noticed in material and method); 100 $\mu\text{L}$ applied on plate (MEA, $37^\circ\text{C}$ ); \*\*micro-pellets of ca 50-200 $\mu\text{m}$ in diam. $\sim 2.0 \times 10^4/\text{mL}$ ) after 10-12 d incubation submerge YES5%, 140 rpm, $25^\circ\text{C}$ ; 100 $\mu\text{L}$ applied on plate (MEA, $37^\circ\text{C}$ ); \*\*\*mycelium without sporulation; colony growing on MEA plate after 4 d incubation ( $37^\circ\text{C}$ ) exposed to high temperature ( $40 \pm 0.5^\circ\text{C}$ )

**FIG 4.**
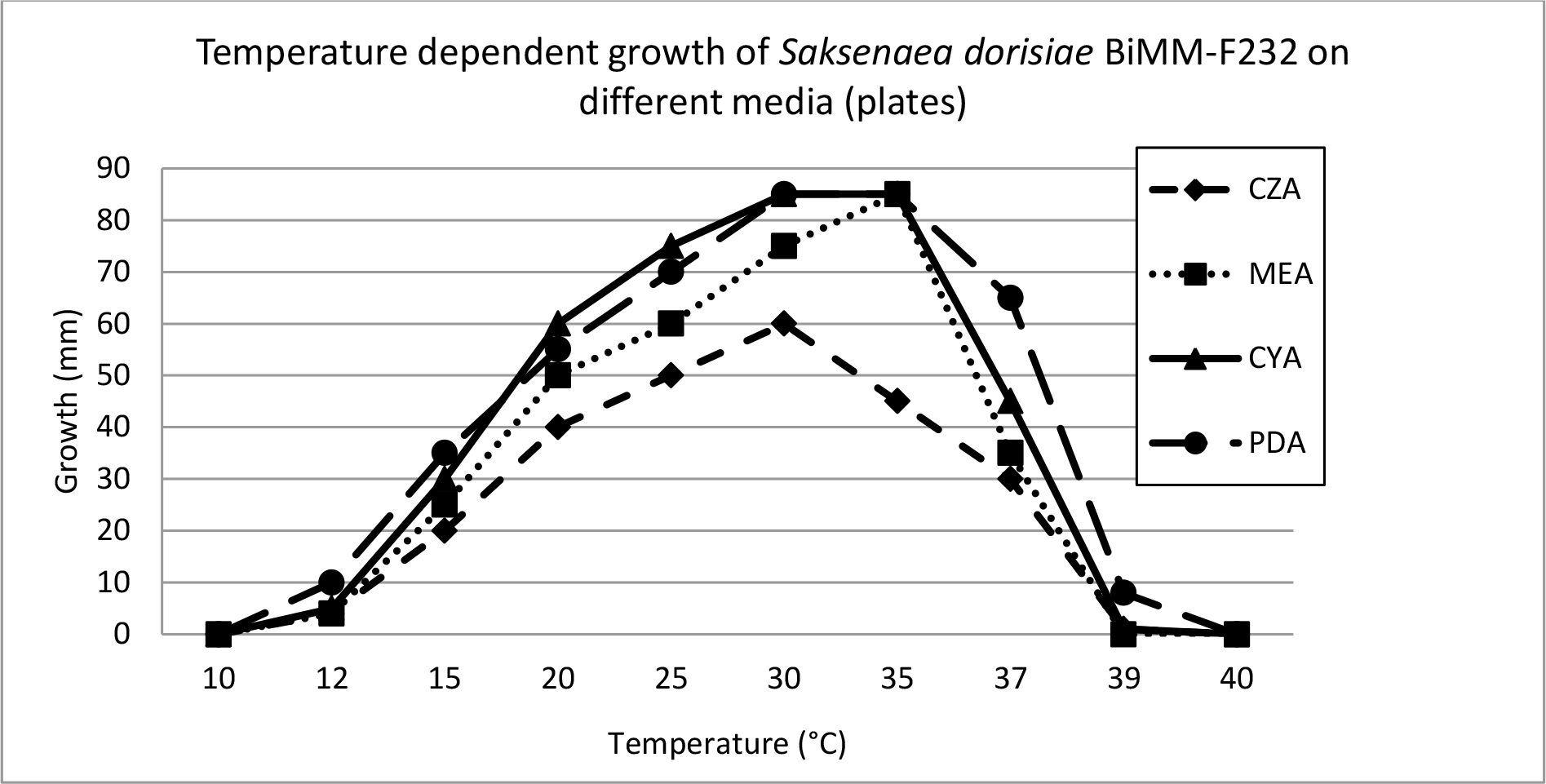
Temperature dependent growth of *Saksenaea dorisiae* (BiMM-F232) and its maximal colony extension (in mm) on Czapek agar (CZA), Malt extract agar (MEA), Czapek yeast extract agar (CYA) and Potato dextrose agar (PDA)

Micromorphology (Fig. 2) – Hyphae mostly coenocytic (non-septate), branched, hyaline, smooth and thin walled, up to 22 µm wide. Sporangiophores erect, generally arising singly, unbranched, brown,, 75-130 long (mostly 85-100 µm), 6-12 µm wide (mostly 7-10 µm), slightly to distinctively verrucose (asperulate) covered with bacilliform protuberances, terminating into hemispherical columellae (15-35 µm wide), and with sparse, dichotomously branched rhizoids (root-like structure). Sporangia terminal, multi-spored, hyaline, flask-shaped (vasiform), slightly to distinctively verrucose (asperulate), 70-190 µm long (mostly 90-160 µm), at maturity with a conical venter up to 55 µm wide (mostly 40-50 µm) and gradually narrowing into a long neck (70-100 µm × 6-10 µm) with a rounded apex (closed with a mucilaginous plug) in young sporangia (Fig. 2 F, G and H), truncated and opened at maturity. Sporangia observed after 5 days on CZA most abundantly at 30°C (up to 30-50 per plate), less so at 25 and 35°C (up to 10 per plate), while none at 20 and 37 °C. The sporangia were formed at the center of the colony nearby or at agar block used for inoculation. Sporangiospores during development and at maturity (4-8 days) mainly short-cylindrical (capsulate) with rounded ends, a few also more or less trapezoidal in lateral view, smooth, thin walled, hyaline, (4.5−)5.0−5.5(−6.0) × (2.0−)2.5−3.0(−3.5) µm (mean = 5.1±0.4 × 2.8±0.3 µm, n = 70). Zygospores not observed.

The main distinguishing phenotypic characteristics of the new species compared with the other taxa of the genus *Saksenaea* are listed in the Table 2.

### Holotype

Serbia, Manastirica (Petrovac) isolated from a private, 65 m deep well – water sample (Code DOO33) 08.10. 2018, isolated by Doris Hochenauer; PRM 951593 [dried culture].

Ex-type strain: BiMM-F232 = CCF 6174.

DNA sequences: GenBank MK559697 (ITS), GenBank MK569515 (TEF-1α), GenBank MK570305 (LSU).

Based on a search of NCBI’s GenBank nucleotide database, the closest hits using the ITS sequence are *S. trapezispora* (UTHSC DI 15-1; Genbank: NR_147690; Identities = 601/644 (93%), gaps 23/644 (3%)) and *S. oblongispora* (CBS 133.90; Genbank: NR_137569; Identities = 595/652 (91%), gaps 35/652 (5%)). By using TEF-1α sequence the closest hits are *S. oblongispora* (CBS 133.90; Genbank: HM776687; Identities = 473/476 (99 %), no gaps) and *S. trapezispora* (UTHSC DI 15-1; Genbank: LT607408; Identities = 467/476 (98%), no gaps)). A region of the 28S rRNA, containing D1 and D2 regions (LSU) shows highest similarities with *S. trapezispora* (GenBank NG_060019; identities = 704/720 (98%), gaps 1/720(0%)) and *S. oblongispora* (GenBank NG_057868; identities = 702/719 (98%), no gaps). The phylogenetic trees built for each ITS (Fig. 5) and LSU (Fig. 6) as well as by combining ITS, LSU and TEF (Fig. 7), indicate that the isolate represents a new species, being closest to *S. trapezispora*.

**FIG 5.**
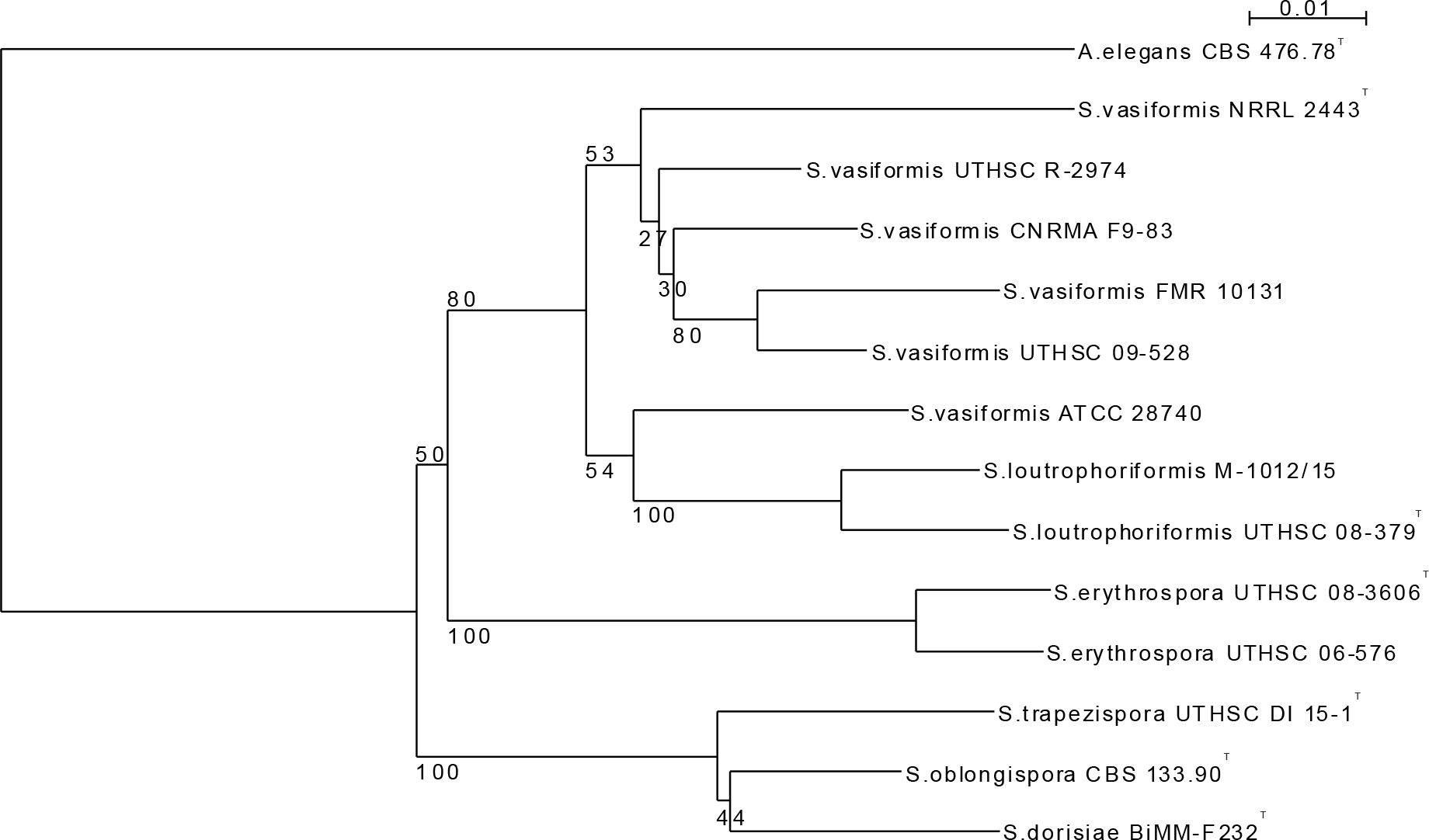
Neighbor joining tree constructed with sequences of the ITS1*-*5.8S-ITS2 of rDNA. The new taxon *S. dorisiae* is compared with species from the same genus. Numbers at nodes indicate bootstrap values (expressed as percentages of 1,000 replications). *Apophysomyces elegans* was used as outgroup. Scale bar indicates 0.01 substitutions per nucleotide position. *S. = Saksenaea*, *A. = Apophysomyces*. ^T^, type strain.

**FIG 6.**
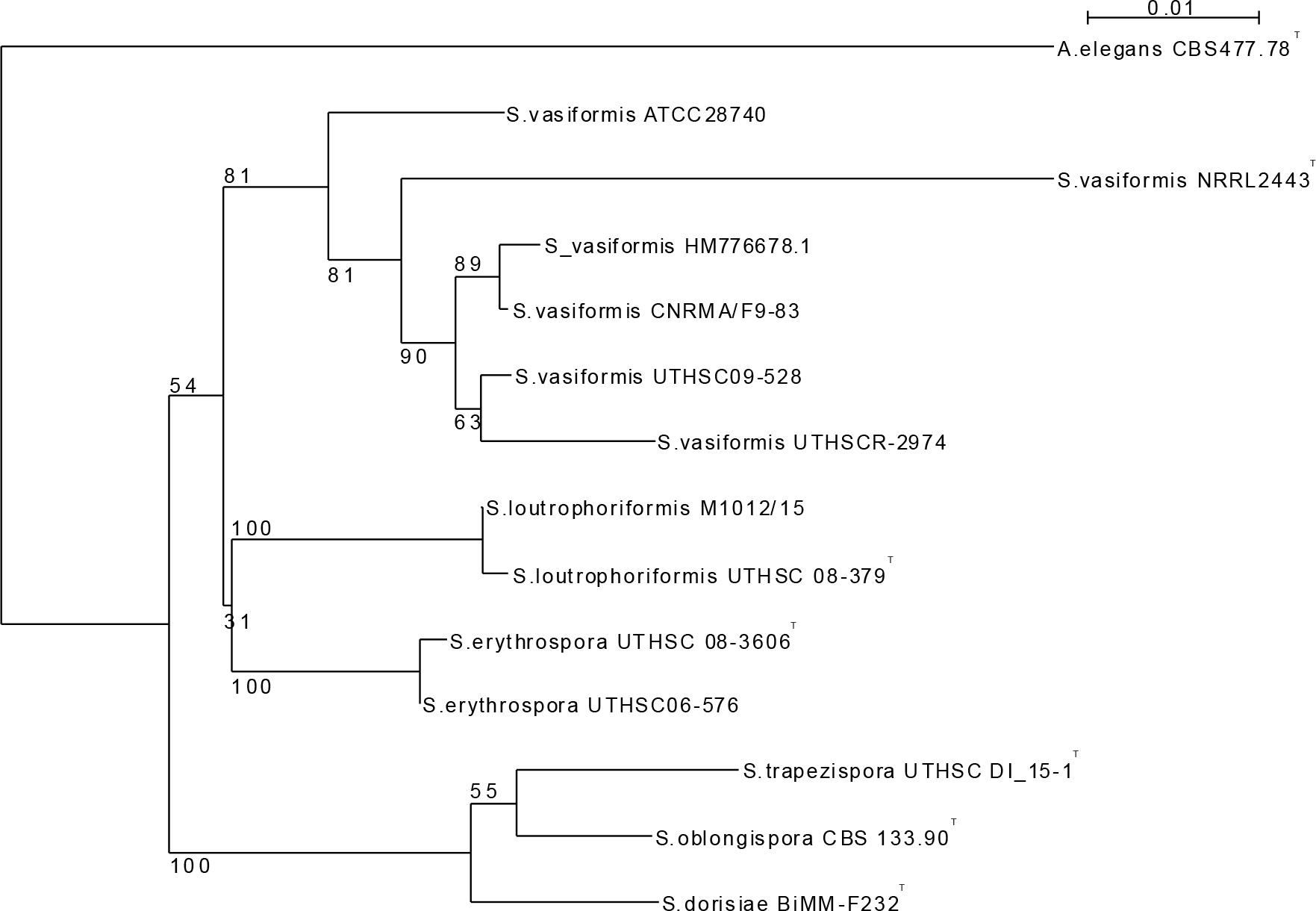
Neighbor joining tree based on a partial sequence the 28S rDNA including D1, D2 domains (LSU) for the new taxon *S. dorisiae* is compared with species from the same genus. Numbers at nodes indicate bootstrap values (expressed as percentages of 1,000 replications). *Apophysomyces elegans* was used as outgroup. Scale bar indicates 0.01 substitutions per nucleotide position. *S. = Saksenaea, A. = Apophysomyces.* ^T^, type strain.

**FIG 7.**
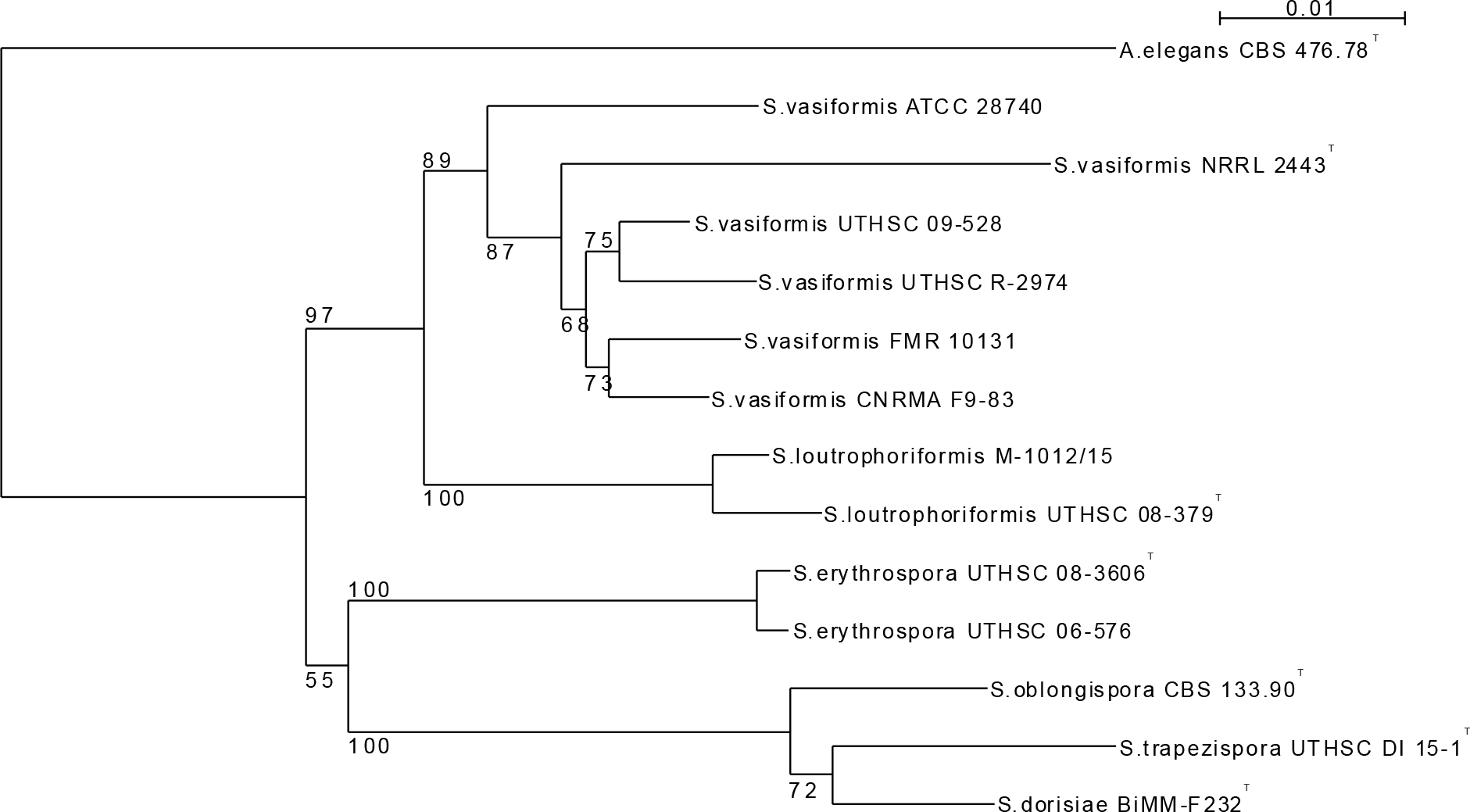
Neighbor joining tree based on a concatenated set of 3 sequences (ITS, LSU and TEF-1α) for the new taxon *S. dorisiae* is compared with species from the same genus *Saksenaea*. Numbers at nodes indicate bootstrap values (expressed as percentages of 1,000 replications). *Apophysomyces elegans* was used as outgroup. Scale bar indicates 0.01 substitutions per nucleotide position. *S. = Saksenaea, A. = Apophysomyces.* ^T^, type strain.

### *In vitro* antifungal susceptibility testing

The MIC Test Strips with 8 antibiotics used for the strain BiMM-F232 (*Saksenaea dorisiae*) sensitivity testing showed that the fungus is sensitive towards ketoconazole, posaconazole and itraconazole at rather high concentrations, i.e. 3.0, 2.0 and 4.0 µg/mL, respectively. The fungus showed resistance towards caspofungin, flucytosine, and voriconazol (MIC > 32 µg/mL) as well as fluconazole (> 256 µg/mL). The MIC value for amphotericin B was unclear as no clear inhibition zone was formed (Fig. 8). Additional application of antibiotic discs (15 antibiotics) with definedconcentrations revealed sensitivity of the strain also towards terbinafine (25 mm zone at 30 µg), ciclopirox (25 mm zone at 50 µg), econazole (15 mm zone at 10 µg) and clotrimazole (25 mm zone at 50 µg). The antibiotic discs containing amphotericin B (with 10 and 20 µg), griseofulvin (10 µg), flucytosin (1 µg), fluconazole (20 and 100 µg), miconazole (10 µg), and voriconazole (1 µg) showed very limited (up to 8mm) or no activity at all (Table 5). It has been observed, however, that all antifungals strips and/or discs, except terbinafine and ciclopirox, were overgrown after incubation for 5 days (37°C). The same type of multi-resistance towards all antifungals (except terbinafine) was observed also when mycelial pellets of the fungus were used for antifungal susceptibility test during this study.

**TABLE 5.**
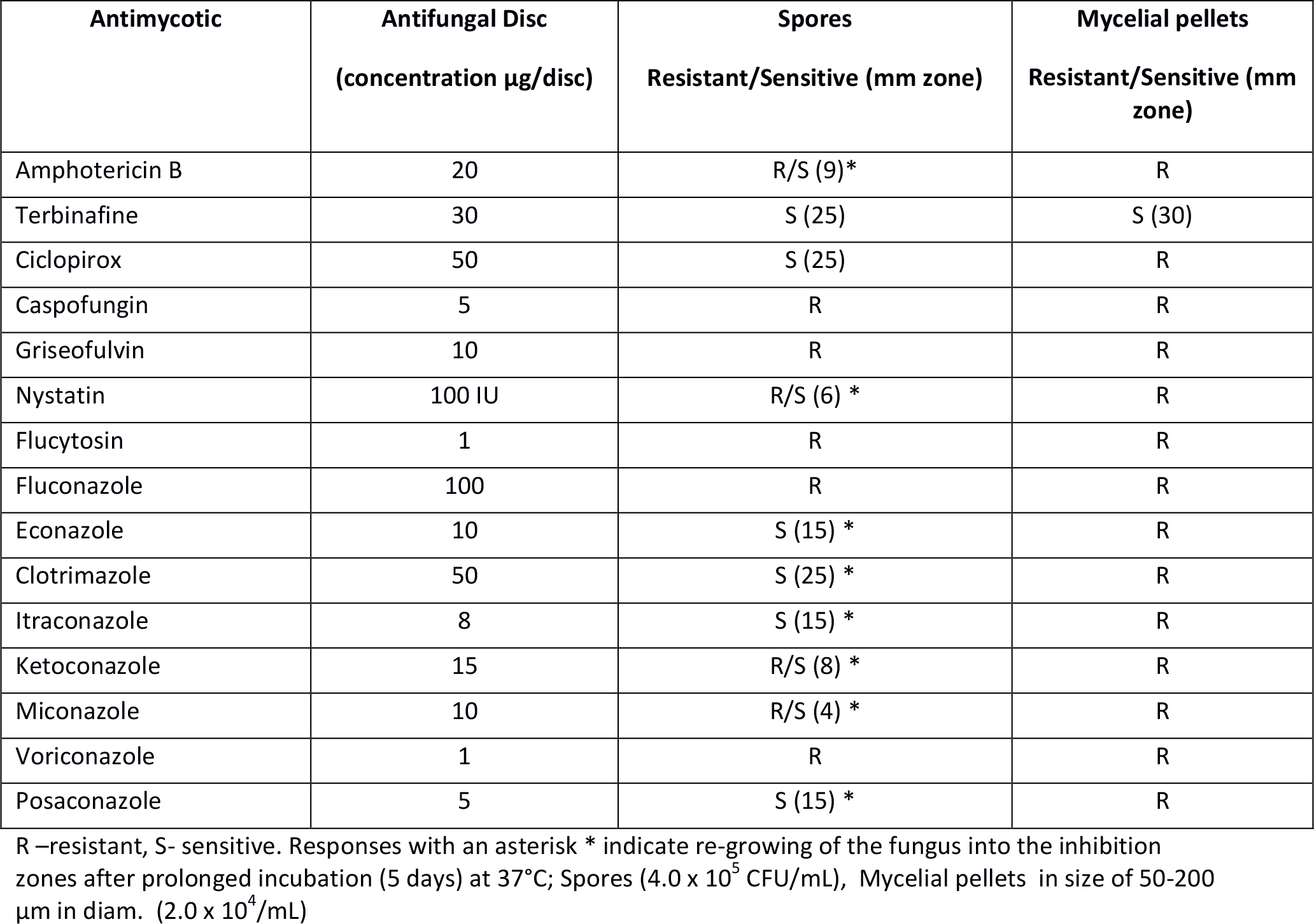
*In vitro* antifungal susceptibility of *Saksenaea dorisiae* (BiMM-F232) towards 15 antifungal discs after 3 days of incubation at 37°C.

**FIG 8.**
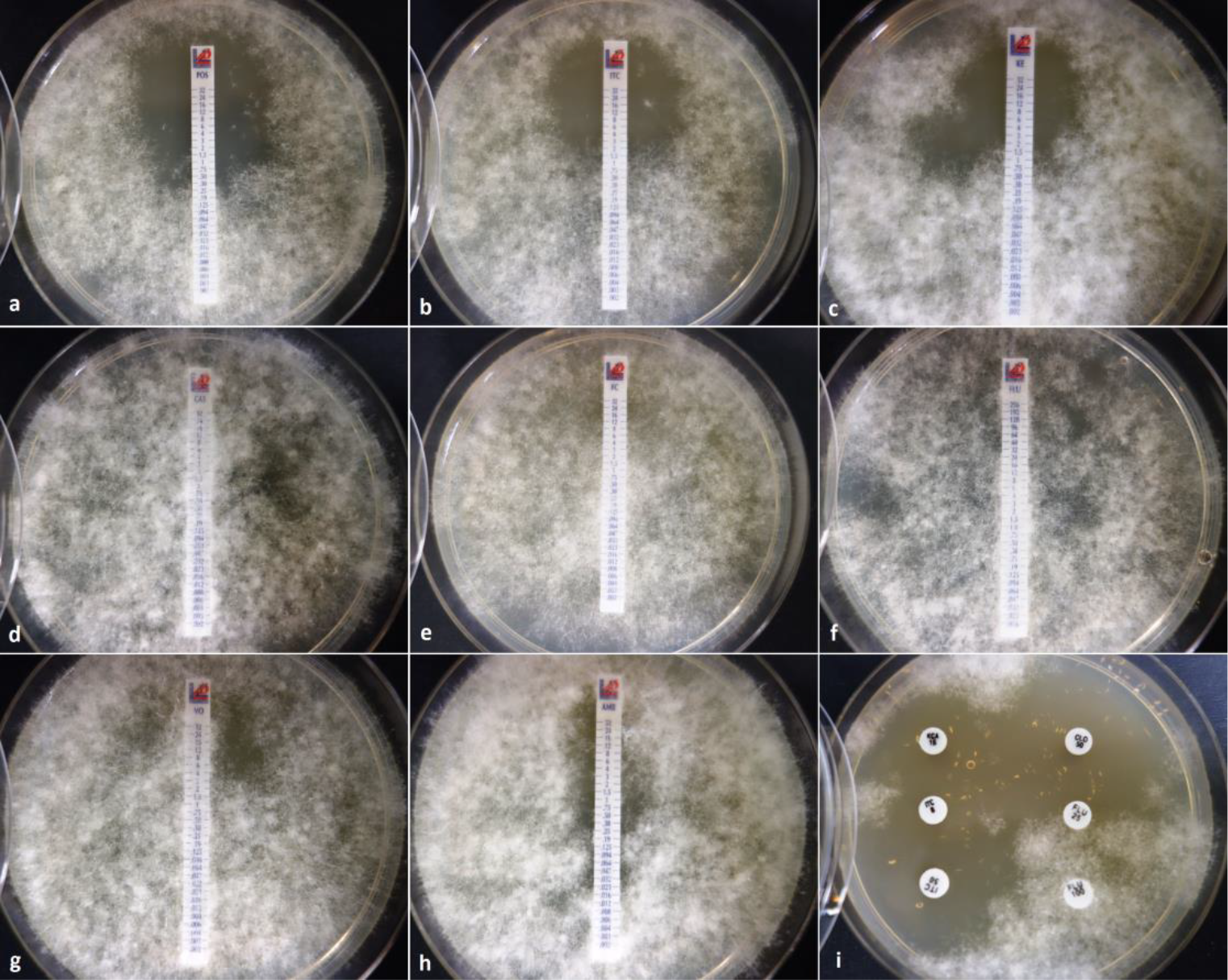
*In vitro* antifungal susceptibility of *Saksenaea dorisiae* BiMM-F232 towards selected antifungals after 3 days incubation at 37°C. MIC Test strips – (a) posaconazole (POS), (b) itraconazole (ITC), (c) ketoconazole (KE), (d) caspofungin (CAS), (e) flucytosine (FC), (f) fluconazole (FLU), (g) voriconazole (VO), (h) amphotericin B (AMB), (i) antifungal discs – left top to left bottom (ketoconazole KCA 15 µg, itraconazole ITC 8 and 15 µg); – right top to right bottom (clotrimazole CLO 50 µm, fluconazole FLU 25 and 100 µm); all plates were overgrown by the fungus after prolonged incubation (5 days) at 37°C.

## DISCUSSION

Phenotypically, the new species, *Saksenaea dorisiae*, is characteristic and differs from the other taxa in the genus *Saksenaea* S.B. Saksena (Mucorales, Saksenaeaceae) by the combination of the following features: (1) no growth at 40°C and slow to moderate growth at 15 and 37°C (15-20 mm and 25-30 mm, respectively), (2) conical sporangial venter, (3) morphology of sporangiospores (short-cylindrical, av. = 5.0 × 3.0 µm), and (4) sparse rhizoids. Especially the shape of sporangia and sporangiospores are very distinctive, rendering *Saksenaea dorisiae* to be easily discernable from the other hitherto known species of the genus. Based on ITS (Fig. 5) and LSU (Fig. 6) *S. dorisiae* is resolved phylogenetically in a cluster with *S. oblongispora* and *S. trapezispora*. Furthermore, phylogenetic reconstruction applying a combination of ITS, LSU and TEF-1α sequences also resulted in clustering of the new species with these two species, being closest to *S. trapezispora* (Fig. 7). All three species do not grow at 42°C in contrast to species in sister clades growing at this temperature, i.e. *S. vasiformis* complex, *S. erythrospora* and *S. loutrophoriformis* (1, 11). At the time of this study, the genus *Saksenaea* contains 6 species, including the new taxon. Each of them is characteristic by a particular combination of morphological traits (mainly morphology of sporangia and sporangiospores) and on this basis also easily distinguished from *S. dorisiae* on these phenotypic bases. The conical-shaped sporangia, sparse rhizoids as well as capsulate sporangiospores formed by the new species are very distinctive, morphologically separating this fungus from all the others in the *Saksenaea* genus. Since it is obvious that formation of sporangia is necessary for phenotypical identification of isolates into the genus and/or species level, incubation on CZA and 30°C are highly recommended conditions to reach sporulation. All species of *Saksenaea* are known for their rapid growth on traditional microbiological media (e.g. SAB, PDA, MEA, WA etc.) but without sporulation. To stimulate sporulation, Padhye et Ajello (17) suggested a simple technique, i.e. floating agar blocks in water with yeast extract, but we could not recapitulate this method with *S. dorisiae*. Out of 9 media used, only CZA was effective in stimulating sporangia development within 4-5 days, at 30°C. This medium has been used for morphological description of all known species of *Saksenaea* (1, 10, 11).

Compared to the phylogenetically close *S. trapezispora* (10), the new species grows substantially slower on CZA at 15°C and 37°C after 4 days (15-20 mm vs 35-45 mm, and 25-30 mm vs 35-40 mm, respectively). Compared to *S. dorisiae*, *S. trapezispora* has longer sporangiophores (up to 230 µm vs 130 µm), spherical instead of conical sporangia, profuse rhizoids versus sparse ones, and substantially larger trapezoid sporangiospores (av. = 7 × 3.5 µm) versus capsulate ones (av. = 5 × 3 µm). In *S. trapezispora*, growth was similar to *S. dorisiae* on CZA at its optimal temperature (30-35°C) after 4 days of incubation. The other species in the genus grow very fast on this medium, their mycelium totally covering the 9 cm Petri dish after 4 days incubation at their optimal temperatures.

Apart of the growth characteristic, shape and size of sporangiospores are important distinguishing morphological characters within the genus (1), also emphasized by Crous et al., (10, 11). In *S. vasiformis* complex and *S. lautrophoriformis* sporangiospores are long cylindrical (to bacilliform) up to 7 × 3.5 µm large, in *S. erythrospora* biconcave (erythrocyte-like) up to 5.5 × 3 µm large, in *S. oblongispora* (ellipsoidal to oblong) up to 6.5 × 4.5 µm large, and in *S. trapezispora* (trapezoidal) up to 8 × 4 µm large. The new species forms mostly short cylindrical (capsulate) sporangiospores, up to 6 × 3 µm large. The phylogenetically closest species, *S. oblongispora* (1), *S. trapezispora* (10), and *S. dorisiae*, are represented by only a single strain. Strain *S. oblongispora* CBS 133.90^T^ has been isolated from a forest soil in Brazil, while a strain *S. trapezispora* CBS 141687^T^ from knee wound of a soldier in Texas, USA.

Mucormycoses caused by *Saksenaea* are difficult to treat. We analyzed sensitivity to antifungal drugs, and found that *S. dorisiae* overgrew all antifungals strips and/or discs used, with exception of terbinafine and ciclopirox. Moreover, the fungus overgrew the inhibition zone after prolonged incubation 5 days at 37°C. Out of 15 antifungals, terbinafine showed stable activity towards the mycelial form (growth from mycelial pellets) of the fungus even after prolonged incubation. It is interesting to note that amphotericin B, a drug that is being the first choice for treatment of mucormycoses (2) including those caused by the *Saksenaea* species (e.g. (6–9), showed only very weak activity against *S. dorisiae*. From the azoles used in the study, ketoconazole, posaconazole and itraconazole showed weak activity at rather high minimum effective concentrations, i.e. 3.0, 2.0, 4.0 µg/mL, respectively. Azoles, such as posaconazole, have been proposed as second-line agents against mucormycosis or as an alternative to amphotericin B in situations when high toxicity of amphotericin B has been detected (2). As indicated here by *in vitro* tests, in case of potential infections caused by *S. dorisiae*, traditional antibiotic therapy applying amphotericin B and posaconazole might not be the optimal treatment strategy. Similarly, in his review on *Saksenaea*, Dannaoui (27) stated that these fungi seem to be less susceptible to amphotericin B than other Zygomycetes. Noteworthy, we found that the fungus is not able to grow at 40°C and is rapidly inactivated at this temperature. After 1 day exposure to 40°C there was no growth or spore germination observed and further cultivation under optimal temperatures demonstrated that the fungus was not surviving this heat treatment. This physiological limitation of the fungus could potentially be used as a subsidiary treatment strategy, along with the suggested terbinafine and/or ciclopirox – therapy.

Even though *S. dorisiae* has been isolated from a single water sample of a private well located in a rural area of the Republic of Serbia (village Manastirica), there is no clear evidence on its connection to water as a primary habitat. Most likely, the fungus originated from surrounding soil resulting in pollution of ground water in the well. In fact, soil is supposed to be the primary habitat of the fungus and a known source for infection typically seen after skin rupture and subsequent contact with contaminated soil (3, 26) or contaminated water (2, 4). Although, the presence of the *Saksenaea* fungi (as *S. vasiformis*) associated with soil environments has been rarely reported, the fungus has been found in different parts of the world, suggesting a wide distribution (27). Species of this genus have been isolated from a clay loam soil in Tamil Nadu state (India), soil from forest tree nurseries in Georgia (the USA), soil in Barro Colorado Island (Panama), in turtle nest sand on the Nancite beach in Costa Rica, soil of banana-producing areas in Cortes province (Honduras), soil of groundnut fields in Israel, soil sample in I-lan prefecture (Taiwan), pineapple field soil in Okinawa (Japan), and also in Africa from intertidal driftwoods Mitsiua (Ethiopia). Infection associated with water or water environment of human and animals (dolphins and a killer whale) have also been reported already (3, 4, 28).

The results of bacteriological investigation (unpublished data) of the water sample revealed a high level contamination by enterococci and other fecal bacterial species indicating that the water of this well was of poor microbiological quality not suitable for consumption. It is noteworthy that the fungus was isolated from an area (village Manastirica, the Republic of Serbia) characterized by cold winters typical for moderately continental climate. Interestingly, we found that the white mycelium of the fungus and mycelial pellets (micro-colonies in a liquid medium) did not survive at 5°C for more than 2 and 7 days, respectively. However, spores in a suspension in water were viable at low temperature for up to 50 days. Thus, especially during cold seasonal periods, spores are vital for propagation and survival of this fungus.

Up to date, all known *Saksenaea* species, were initially described from areas outside of Europe, namely *S. vasiformis* (India: NRRL 2443^T^), *S. lautrophoriformis* (USA: UTHSC 08-379^T^ and India: M-1012/15), *S. erythrospora* (USA: UTHSC 08-3606^T^ and Middle East: UTHSC 06-576), *S. oblongispora* (Brazil: CBS 133.90^T^), and *S. trapezispora* (USA: CBS 141687^T^). To the best of our knowledge, reports dealing with *Saksenaea* fungi from Europe have been associated only with *S. vasiformis* and areimited to the Mediterranean climate area of Spain (3, 13–15) and/or Greece (2), and. Hence, *S. dorisiae* represents a first described species of this genus originating from a moderately continental climate area in Europe.

## ACKNOWLEDGMENTS

The authors thank Stephan Bruck (AQA GmbH, Vienna, Austria) for providing data on the sample origin. The Bioactive Microbial Metabolites research platform (BiMM) is supported by grants K3-G-2/026-2013 and COMBIS/ LS16005 funded by the Lower Austria Science and Education Fund (NfB).

